# Transcriptome analyses of 7-day-old zebrafish larvae possessing a familial Alzheimer’s disease-like mutation in *psen1* indicate effects on oxidative phosphorylation, mcm functions, and iron homeostasis

**DOI:** 10.1101/2020.05.03.075424

**Authors:** Yang Dong, Morgan Newman, Stephen Pederson, Nhi Hin, Michael Lardelli

## Abstract

Early-onset familial Alzheimer’s disease (EOfAD) is promoted by dominant mutations, enabling the study of Alzheimer’s disease (AD) pathogenic mechanisms through generation of EOfAD-like mutations in animal models. In a previous study, we generated an EOfAD-like mutation, *psen1*^*Q96_K97del*^, in zebrafish and performed a transcriptome analysis comparing entire brains from 6-month-old wild type and heterozygous mutant fish. We identified predicted effects on mitochondrial function and endolysosomal acidification. Here we aimed to determine whether similar effects occur in 7 day post fertilization (dpf) zebrafish larvae that might be exploited in screening of chemical libraries to find ameliorative drugs. We generated clutches of wild type and heterozygous *psen1*^*Q96_K97del*^ 7 dpf larvae using a paired-mating strategy to reduce extraneous genetic variation before performing a comparative transcriptome analysis. We identified 228 differentially expressed genes and performed Goseq analysis and gene set enrichment analysis (GSEA). This predicted a significant effect on oxidative phosphorylation, consistent with our earlier observations of predicted effects on ATP synthesis in adult heterozygous *psen1*^*Q96_K97del*^ brains. The dysregulation of minichromosome maintenance protein complex (MCM) genes strongly contributed to predicted effects on DNA replication and the cell cycle and may explain earlier observations of genome instability due to *PSEN1* mutation. The upregulation of *crystallin* gene expression may be a response to defective activity of mutant Psen1 protein in endolysosomal acidification. Extracellular matrix (ECM) related genes were downregulated, consistent with previous studies of EOfAD mutant iPSC neurons and postmortem late onset AD brains. Also, changes in expression of genes controlling iron ion transport were observed without identifiable changes in the prevalence of transcripts containing iron responsive elements (IREs) in their 3’ untranslated regions. These changes may, therefore, predispose to the apparent iron dyshomeostasis previously observed in 6-month-old heterozygous *psen1*^*Q96_K97del*^ EOfAD-like mutant brains.

## Introduction

Alzheimer’s disease (AD) is a progressive neurodegenerative brain disorder that eventually develops into dementia. AD is a serious worldwide health issue and shows a trend of increasing disease incidence [1]. AD may be classified in numerous ways. Late onset, sporadic AD, occurs after 65 years of age and is the most common form, contributing to more than 95% of AD cases [2]. This form of AD is affected by multiple factors, including age, diet, life style, genetic and environmental factors [3]. Therefore, it has been difficult to model in animals. An early onset, familial form of AD (EOfAD) shows autosomal, dominant inheritance and contributes less than 5% of all AD cases [4]. As both AD forms share similar pathologies [2], many researchers model EOfAD through genetic manipulation of animals to study AD ontology and pathology in general.

Rodent models are the most commonly used in AD research. However, current transgenic rodent models used in EOfAD studies do not reflect closely the disease state of human patients. In 2017, Hargis and Blalock [5] summarized brain transcriptional profiles in human AD, and compared five transgenic mouse models of AD to human AD profiles. All of these mouse models failed to model the most consistent transcriptional signature of human AD, a downregulation of neuronal and mitochondrial genes. Also, the focus of most AD studies is on the pathologies of the advanced disease, such as the accumulation of amyloid-β peptide and tau protein, and on identification of new biomarkers for early diagnosis. Our laboratory seeks deeper insight into the early molecular states of AD brains to explore disease etiology and molecular mechanisms. We have modeled EOfAD-like mutations in another popular vertebrate animal model, the zebrafish. The zebrafish has a fully sequenced and well annotated genome [6], and has the advantages of rapid development with a relatively short generation time. It is easily manipulated genetically and has the capacity to produce large families of siblings which can then be raised together in the same environment to limit the effects of environmental and genetic noise in molecular analyses [7]. Moreover, zebrafish possess orthologs of the human genes mutated in EOfAD. Most recognized EOfAD-causative mutations have been found in the genes *PSEN1, PSEN2* and *APP* [8]. (The majority of these mutations, ∼63%, occur in the gene *PSEN1* [9].) The zebrafish orthologs of these genes have been identified as *psen1* [10], *psen2* [11], *appa* and *appb* [12]. Therefore, zebrafish have the potential to model EOfAD mutations for the study of the molecular pathological processes of AD. The zebrafish is also a versatile model for drug screening as its tiny larvae can be obtained in large numbers and arrayed into microtitre plates for molecular, developmental, or behavioural analyses [13].

One EOfAD-like mutation we have generated is *psen1*^*Q96_K97del*^, a deletion of 6 nucleotides in the zebrafish *psen1* gene. This mutation deletes 2 codons but maintains the open reading frame, leading to structural and hydrophilicity changes in the first lumenal loop of the translated protein. Although this mutation is not the exact equivalent of any currently known human EOfAD mutation, there are numerous similar EOfAD mutations that distort the first luminal loop of human *PSEN1* (e.g. *PSEN1*^*L113_I114insT*^ [14], *PSEN1*^*P117L*^ [15]) and, like all the many various and widely distributed EOfAD mutations in the *PRESENILIN* genes, it follows the “fAD mutation reading frame preservation rule” [8].

Like human EOfAD mutations, *psen1*^*Q96_K97del*^ has dominant effects when heterozygous. We have observed that the brains of 6-month-old (young, recently sexually mature adult) zebrafish heterozygous for *psen1*^*Q96_K97del*^ show transcriptome alterations consistent with disturbances in energy production (ATP synthesis) and lysosomal dysfunction [16]. These may represent the initial stresses that, after decades in humans, lead to AD.

Could our heterozygous mutant zebrafish be used to identify drugs that suppress these molecular defects and so prevent the pathological progression to AD? A 2015 paper by Wagner et al. [17] showed that the most effective drugs in an animal model (of dyslipidemia) were those that best caused reversion of the transcriptomic disease signature to normal. In accordance with this philosophy, we might use our zebrafish mutants to screen for AD-preventative drugs based on the drugs’ ability to revert transcriptomic signatures of ATP synthesis disruption and lysosomal dysfunction back to wild type. Therefore, as a first step in assessing the viability of this idea, we were interested to observe whether the transcriptomic signatures evident in 6-month-old zebrafish *psen1*^*Q96_K97del*^ heterozygous adult mutant brains were discernable in whole zebrafish larvae.

Our previous analysis of *psen1*^*Q96_K97del*^ heterozygous adult mutant brain transcriptomes was facilitated by the ability to perform bulk RNA-seq on the entire ∼7 mg brains of individual mutant zebrafish and their wild type siblings. While an individual zebrafish larva at 7 days post fertilization (dpf, when feeding would normally begin) is too small to provide sufficient RNA for bulk RNA-seq analysis without some form of amplification, we can produce clutches of uniformly heterozygous larvae by crossing a homozygous mutant parent fish with a wild type parent. Analysis of pooled RNA from multiple individuals also reduces between-genotype variability due to “averaging” of the mRNA expression levels contributed by each larva in the pool. Also, using a single male fish to produce both a heterozygous mutant clutch and a wild type clutch of larvae (though mating with a single homozygous mutant or wild type female fish respectively) further reduces genetic variability in the analysis (see Figure 1).

**Figure 1.**
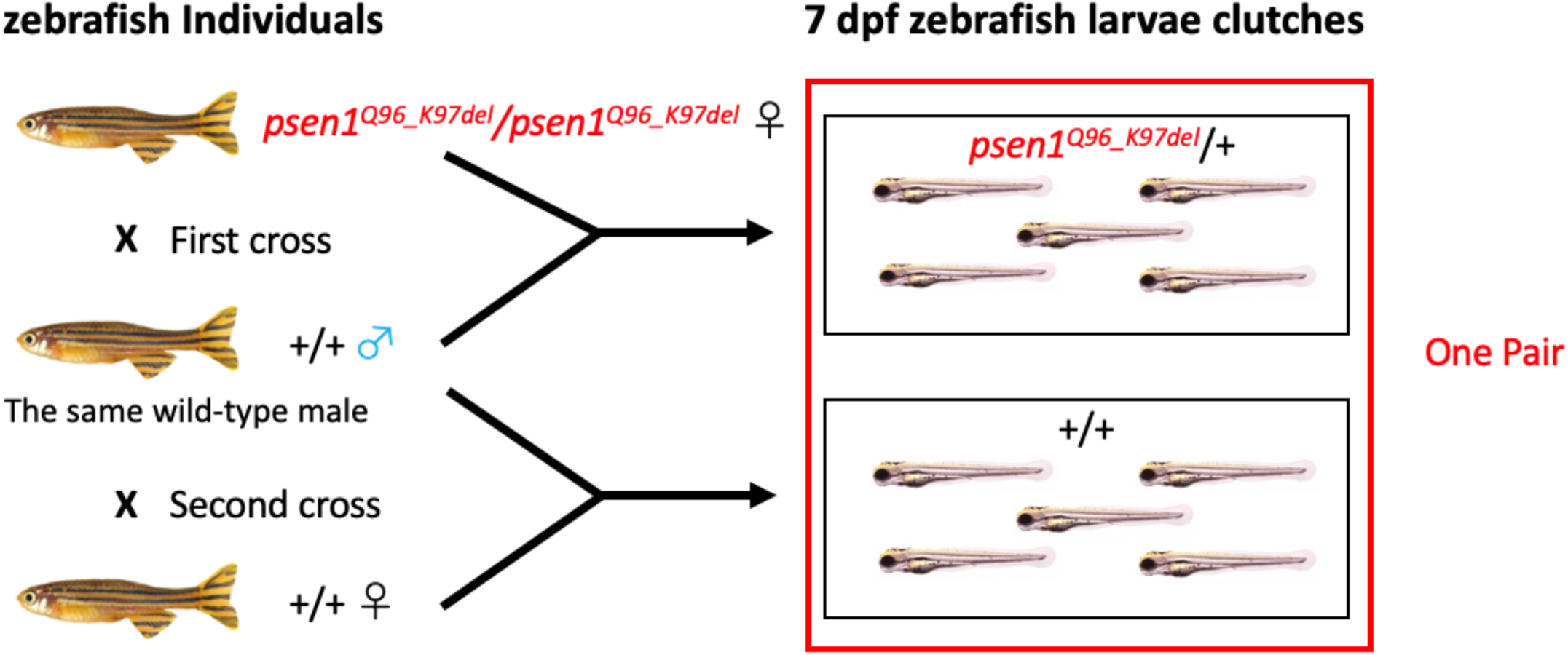
Mating scheme to generate pairs of 7 dpf zebrafish larval clutches.

In this paper we describe a transcriptome analysis on clutches of 7 dpf heterozygous mutant and wild type larvae structured as described above to minimize genetic variation. This identified 228 potentially differentially expressed (DE) genes. Bioinformatic predictive analysis identified probable significant changes in DNA replication and cell cycle processes, to which changes in the regulation of genes related to the minichromosome maintenance protein complex (MCM) were the main contributors. In addition, effects on iron ion transport were identified, suggesting a potential early disruption of iron homeostasis components that might lead, ultimately, to mitochondrial dysfunction including disruption of ATP synthesis.

## Materials and Methods

### Ethics

The research described in this paper was carried out under permit S-2017-073 from the Animal Ethics Committee of the University of Adelaide.

### Zebrafish pair-mating breeding strategy

A female zebrafish homozygous for the *psen1*^*Q96_K97del*^ allele was crossed to a male wild-type fish to generate a clutch of heterozygous *psen1*^*Q96_K97del*^/+ larvae. The same male wild-type fish was then crossed to a female wild-type fish to generate a group of wild-type embryos (Figure 1). The use of the male fish as the common parent in this mating scheme avoids the possibility that maternal effects might obscure transcriptome differences between the larval genotypes [18]. These two clutches of larvae from an individual male wild-type fish were labeled a “pair” (Figure 1). For statistical power, we analysed n=6 pairs of larval clutches. Note that no individual adult fish contributed to more than one larval clutch pair.

### Sample preparation and RNA purification

Six pairs of larval clutches were generated as described above (12 clutches in total). The parents of 4 pairs of clutches were approximately 5 months old, while the parents of the other two pairs of clutches were approximately 12 months old. The larvae were raised in E3 medium for 7 days, allowing complete larval yolk absorption but not food intake. This was done to minimize the influence of environmental factors on gene expression. As individual larvae are too small to generate sufficient RNA for bulk RNA-seq analysis, 40 larvae were pooled from each clutch.

Total RNA was extracted from pooled larvae using the mirVanaTM miRNA isolation Kit (Ambion, Life Technologies, Thermo Fisher Scientific, Waltham, MA, USA). DNase treatment was performed on RNA to remove remaining genomic DNA using the DNA-free™ Kit (Ambion, Life Technologies, Thermo Fisher Scientific, Waltham, MA, USA). Purified total RNA was then delivered to the Genomics Facility at the South Australian Health and Medical Research Institute (SAHMRI, Adelaide, Australia) for RNA sequencing. Demultiplexed libraries were provided by the sequencing centre at SAHMRI as 75bp single-end reads, after using polyA amplification. All libraries were sequenced to a minimum depth of 30 million reads, across two NextSeq lanes which were subsequently merged. The data has been deposited in NCBI Gene Expression Omnibus (GEO) database [19] and are accessible through GEO accession number GSE148631.

### Transcriptome data treatment and DE gene identification

Reads from each sample were trimmed using AdapterRemoval [20] setting a minimum quality of 30 and a minimum length of 35. Trimmed reads were aligned to the *Danio rerio* genome Ensembl Release 96 (GRCz11) using STAR v2.5.3a [21], and the aligned reads were assigned to each gene using featureCount [22], specifying that only unique alignments which strictly overlapped exonic regions were counted. The 19,396 genes that received more than 1 count per million reads (CPM) in at least 6 samples, were retained for further analysis. The remaining 12,661 genes were discarded as undetectable genes, giving library sizes ranging between 44,498,373 and 52,779,608 reads. A design matrix was specified with an intercept for each pair and with genotype as the common difference. An initial Differential Gene Expression (DE) analysis was then performed using the glmLRT method as implemented in edgeR [23, 24]. The 5000 lowest ranked genes from this analysis (i.e. with the highest p-value) were selected as unchanged negative control genes and passed to RUVg [25], setting k = 1. After RUV treatment, the same design matrix with the addition of the offset term from RUV was used to perform a new DE analysis. An FDR cutoff of 0.01 was then used to identify a gene as differentially expressed (DE).

### Goseq analysis and Gene set enrichment analysis (GSEA)

The mappings linking zebrafish genes to Hallmark/KEGG pathways, Wiki pathways and GO terms were achieved by msigdbr [26], rWikipathways [27] and org.Dr.eg [28] respectively. To look for enrichment of gene-sets within our defined set of DE genes, a Probability Weighting Function (PWF) was calculated based on the set DE genes using gene lengths as the bias data. Then goseq analysis [29] was performed using this PWF and gene-sets were considered as significantly enriched in the set of DE genes using a FDR threshold of 0.05.

In order to look at the more complete list of all expressed genes, we then used Gene Set Enrichment Analysis (GSEA, [30]) as implemented in the fgsea [31] package. A ranked list was formed using sign(log_2_FC) * (-log_10_PValue) as the ranking statistic and the same gene-sets as for goseq were tested, using n = 10^5^ iterations.

A Bonferroni corrected p-value < 0.05 was used to identify significantly-altered pathways. Pathway Diagrams for detected KEGG pathways were plotted by pathview [32] using log_2_FC.

### Iron responsive elements (IRE) enrichment analysis

Genes containing iron responsive elements (IRE) in either their 5’ or 3’ untranslated regions (UTRs) were identified using searching for IREs (SIREs) [33]. Both goseq [29] and GSEA were performed to detect the enrichments of the genes containing IREs in their 5’UTR or 3’UTR respectively. An FDR and a Bonferroni corrected p-value < 0.05 were used to identify significant enrichments in goseq analysis and GSEA respectively.

## Results

Our previous study examined the effects of heterozygosity for the *psen1*^*Q96_K97del*^ mutation on the transcriptome of 6-month-old zebrafish brains. The changes in gene expression observed were predicted to affect ATP synthesis and lysosomal acidification [16]. Here we sought to identify the changes present in entire, heterozygous 7 dpf larvae to assess whether these larvae might be a suitable system in which to screen drug libraries for compounds ameliorating the young adult brain ATP synthesis and lysosomal acidification effects. The mating scheme described in Figure 1 was employed to generate n=6 pairs of heterozygous mutant and wild type clutches of larvae. (Power calculations performed since our first publication indicated that n=6 provides a power of approximately 70% for detection of fold-change > 2 at a false discovery rate of 0.05 across the vast majority of expressed transcripts in zebrafish brain transcriptomes [34], data not shown.) RNA-seq was performed on RNA purified from these clutches followed by a comparative transcriptome analysis to identify differentially expressed genes and explore potential functional effects caused by the mutation.

### Differentially expressed genes (DE genes)

Gene expression differences between wild type and heterozygous *psen1*^*Q96_K97del*^/+ clutches were calculated through a design matrix considering each pair of clutches as a factor and genotypes as the common difference.

228 significantly DE genes were identified (Supplementary data 1), although most of these genes showed only minor fold-change differences in expression. Comparison of the significantly DE genes identified from heterozygous mutant 7 dpf larvae with those seen in heterozygous mutant 6-month brains [16], revealed only one gene, *lgals8b*, as common between the two datasets. It is upregulated in both.

To support the accuracy and reliability of the RNA-seq data, relative standard curve quantitative PCRs (qPCRs) were performed for four of the most statistically significantly DE genes that showed relatively large fold-changes in expression. The qPCRs were performed using cDNA synthesized from the same preparations of RNA that were used in the RNA-seq analysis. Three of the four genes were seen to be differentially expressed to a statistically significant degree (p<0.05, Supplementary data 2).

### GOseq analysis of pathways and GO terms

To predict the cellular functions affected by heterozygosity for the *psen1*^*Q96_K97del*^ mutation, we analysed the DE genes using the Hallmark, KEGG, and Wiki pathway databases and the Gene Ontology database. Different pathway databases may contain different representations of similar biological pathways. Hallmark gene sets summarize well-defined biological states or processes built on the overlapping of several gene set collections, and so are useful to achieve an overall view [26]. The KEGG and Wiki gene sets are two popular pathway databases allowing examination of high-level functions. Different pathway databases might show low between-database consistency due to the incomprehensive gene sets and gene interactions in each category [35]. Therefore, to generate a more comprehensive result, we used both KEGG and Wiki pathway databases for pathway analysis.

Pathway and GO analysis were performed using Goseq, which weighted DE genes and calculated each category’s significance amongst DE genes to identify significantly changed pathways or GO terms. Goseq analysis only focuses on the proportions of DE genes in each category but does not consider gene expression fold change and pathway regulation direction. Table 1 shows the Goseq results with an FDR cutoff of 0.05 in the analysis of Hallmark, KEGG and Wiki pathways (Table 1) and of GO terms (Figure 2). In the Hallmark pathway (Table 1), G2M_CHECKPOINT contains genes critical for cell division cycle progression, and E2F_TARGETS includes numerous genes that play essential rolls in the cell cycle and DNA replication [36]. Therefore, the Goseq results of the Hallmark, KEGG and Wiki pathway analyses (Table 1) show significant changes in DNA replication and cell cycle control. Among the DE genes in these two categories, most are members of the minichromosome maintenance (MCM) protein family. Downregulation of the genes *mcm2, mcm3, mcm4, mcm5, mcm6* and *mcmbp* and upregulation of the gene *mcm7* were observed in the heterozygous mutant larvae.

**Table 1.**
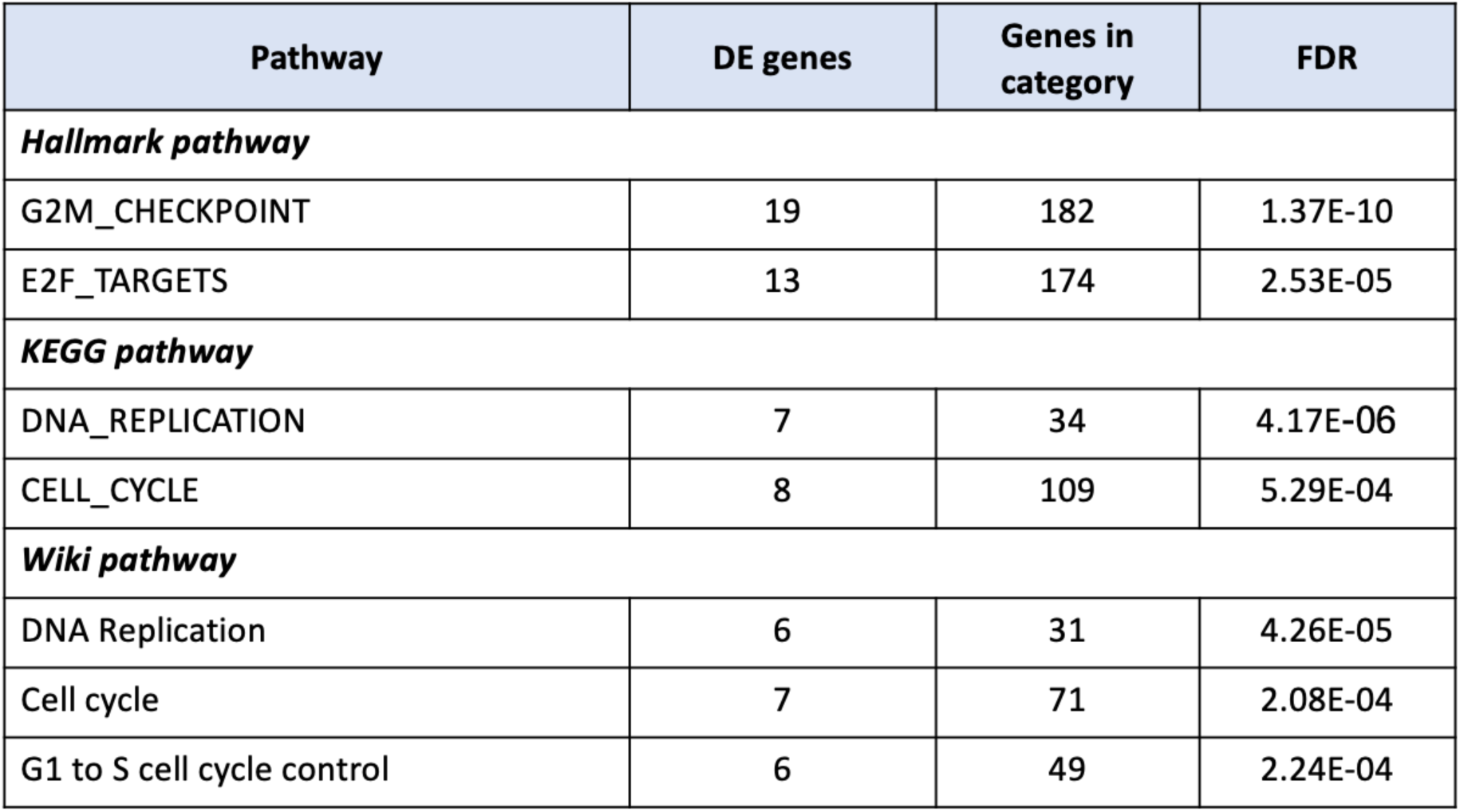
Significantly-changed pathways in the Goseq analysis of Hallmark, KEGG and Wiki pathways filtered by a FDR cutoff of 0.05.

**Figure 2.**
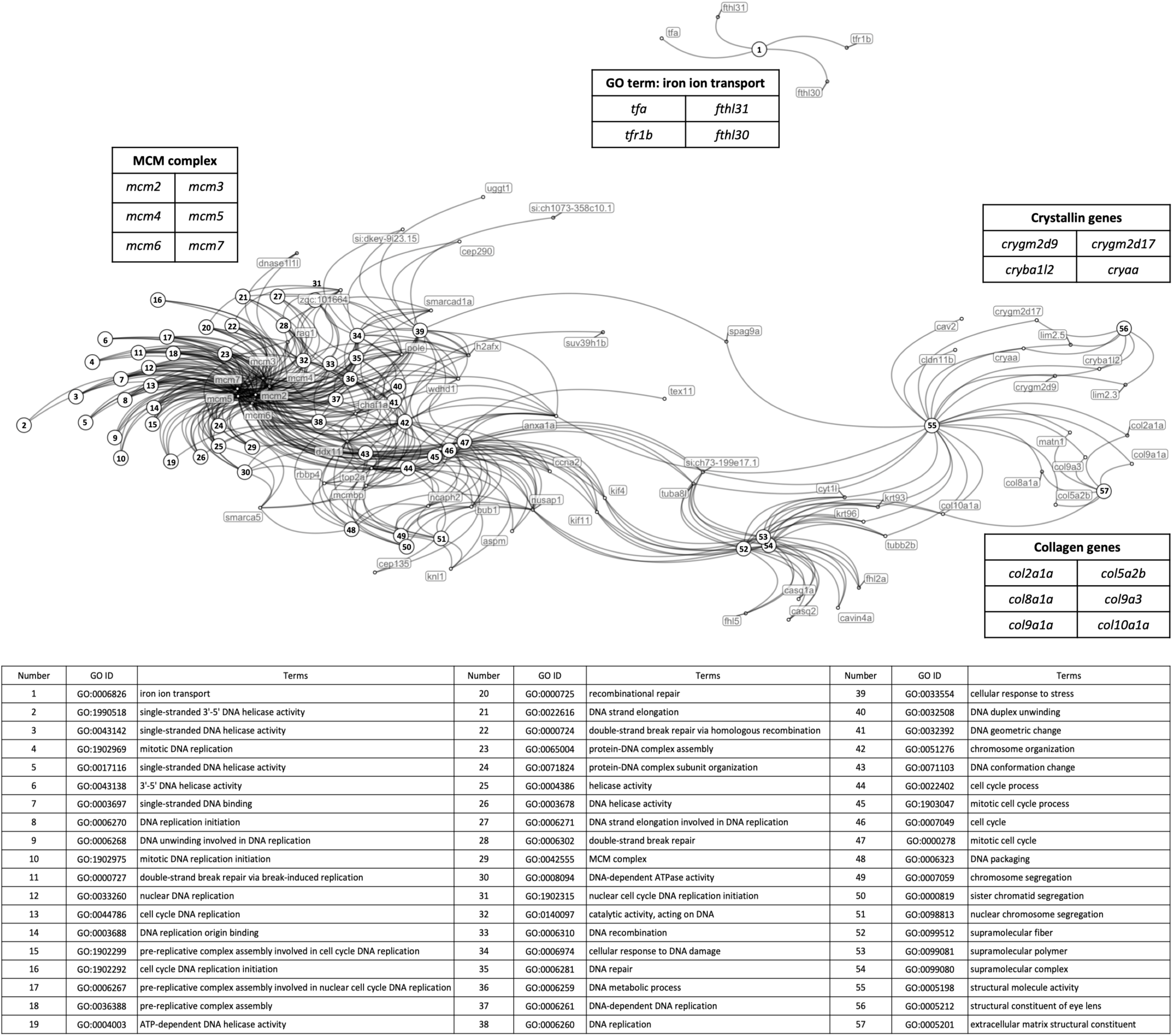
Network of relationships between DE genes and significantly-changed GO terms in the Goseq analysis. Dots represent DE genes and are labelled with gene names., Numbered circles represent those GO terms showing significant enrichment for the DE genes. The table below the network indicates the GO represented by each number.

In GO analysis, one DE gene can contribute to several related GO terms. The network shown in Figure 2 illustrates how the DE genes are shared between GO terms. Similar to the pathway analyses, most of the GO terms showing significant enrichment for DE genes are related to the cell cycle and DNA replication. In the network, these GOs cluster around the *MINICHROMOSOME MAINTENANCE (mcm)* genes. The network also illustrates how numerous genes can form a functionally related cluster contributing to only one or a few GOs. This is seen for the significantly upregulated *CRYSTALLIN* genes that contribute to eye lens structure (GO: *Structural constituent of eye lens*) but also function in lysosomal acidification (not revealed here, see Discussion). In contrast, the four genes included in the GO *Iron ion transport* show significantly changed regulation. This includes downregulation of the genes *tfa* and *tfr1b* that act to import iron via the endolysosomal pathway [37]. The *ferritin heavy chain like* genes *fthl30* and *fthl31* are upregulated and downregulated respectively, presumably influencing the storage of ferric iron within cells.

We recently published an analysis using a novel method of transcriptome analysis to detect differences in ferrous iron (Fe^2+^) status in cells [38]. Using this technique, we detected for the first time, that young (6-month-old) adult brains from *psen1*^*Q96_K97del*^/+ zebrafish are likely deficient for ferrous iron. Therefore, we were very interested to see evidence of iron ion transport gene expression changes in the 7 dpf *psen1*^*Q96_K97del*^/+ larvae. To confirm the reality of this changed gene expression we performed qPCRs for the genes *tfa, tfr1b*, and *fthl31* on cDNA made from the same mRNA samples that were subjected to RNA-seq (see Supplementary data 2, *fthl31* was not examined because its expression level is particularly low). The qPCRs for these three genes were consistent with the RNA-seq results.

When ferrous iron is deficient in cells, Iron Regulatory Proteins bind to Iron-Responsive elements in the 3’ untranslated regions (3’UTRs) of mRNAs encoding proteins that function to increase ferrous iron levels (such as human TFR1 [39] or zebrafish Tfr1b [40]). To detect ferrous iron dyshomeostasis in transcriptome data, we looked for enrichment of a large set of gene mRNAs that include putative IREs in their 3’ UTRs. We did not see enrichment of this gene set in the 7 dpf *psen1*^*Q96_K97del*^/+ zebrafish larvae, likely indicating that the apparent ferrous iron deficiency of young adult *psen1*^*Q96_K97del*^/+ brains requires time to develop (Supplementary data 4).

### Gene set enrichment analysis (GSEA)

Goseq analysis only focuses on significantly DE genes and predicts affected pathways based on DE gene numbers in each GO. In contrast, GSEA ranks all genes based on fold change and P-value, and then estimates their contributions to each pathway. Therefore, GSEA can show pathway regulation direction, and provides a complementary view of gene sets.

We applied GSEA using the Hallmark, KEGG and Wiki pathway databases. Several significantly-changed pathways were identified in each analysis (Table 2), including pathways previously identified in the Goseq pathway analysis. Four of the significantly-changed KEGG pathways are illustrated in Figure 3. *DNA replication* (Figure 3a) and *cell cycle* (Figure 3b) were the most significantly affected pathways identified in the Goseq pathway analysis and the GO analysis. Regulation of the MCM complex plays essential roles in both pathways. The MCM complex forms a DNA helicase, which cooperates with replication protein A (RPA) to unwind duplex parental DNA before DNA synthesis (Figure 3a, [41]). Dysregulation of the MCM complex would influence the DNA replication and might cause replication stress leading to genomic instability [42]. The pathways *ECM receptor interaction* (Figure 3c) and *oxidative phosphorylation* (OXPHOS, Figure 3d) were also significantly changed in 7 dpf *psen1*^*Q96_K97del*^/+ zebrafish larvae. *ECM receptor interaction* was the most significantly changed pathway in KEGG pathway analysis (the lowest P-value), and most genes involved were downregulated (Figure 3c), including the *COLLAGEN* gene group identified in the previous GO analysis. The KEGG pathway *ECM receptor interaction* plays important roles in control of cellular activities, including functioning to provide cell structural support and to regulate cell-cell and cell matrix interactions [43]. In developing brains, *ECM receptor interaction* participates in cell migration and the guidance of growing axons, having crucial effects on neural cells. This has implicated *ECM receptor interaction* in processes underlying many central nervous system diseases such as AD, schizophrenia and Parkinson’s disease [44]. OXPHOS (Figure 3d), as well as *fatty acid metabolism* (shown in Table 2), contribute to the fundamentally important function of energy production. In our previous GO analysis of 6-month-old *psen1*^*Q96_K97del*^/+ zebrafish brains, we saw very significant apparent effects on ATP synthesis [16]. The analysis here suggests that that energy production capacity is downregulated in the mutant larvae and this is expected to include ATP synthesis. Furthermore, *Beta-alanine metabolism, glutathione metabolism, pyrimidine metabolism, butanoate metabolism* and *focal adhesion* are also identified as significantly-changed pathways (Table 2). The interpretation of these pathway changes requires further investigation. KEGG diagrams for the statistically significantly affected pathways not shown in Figure 3 are given in Supplementary data 3.

**Table 2.**
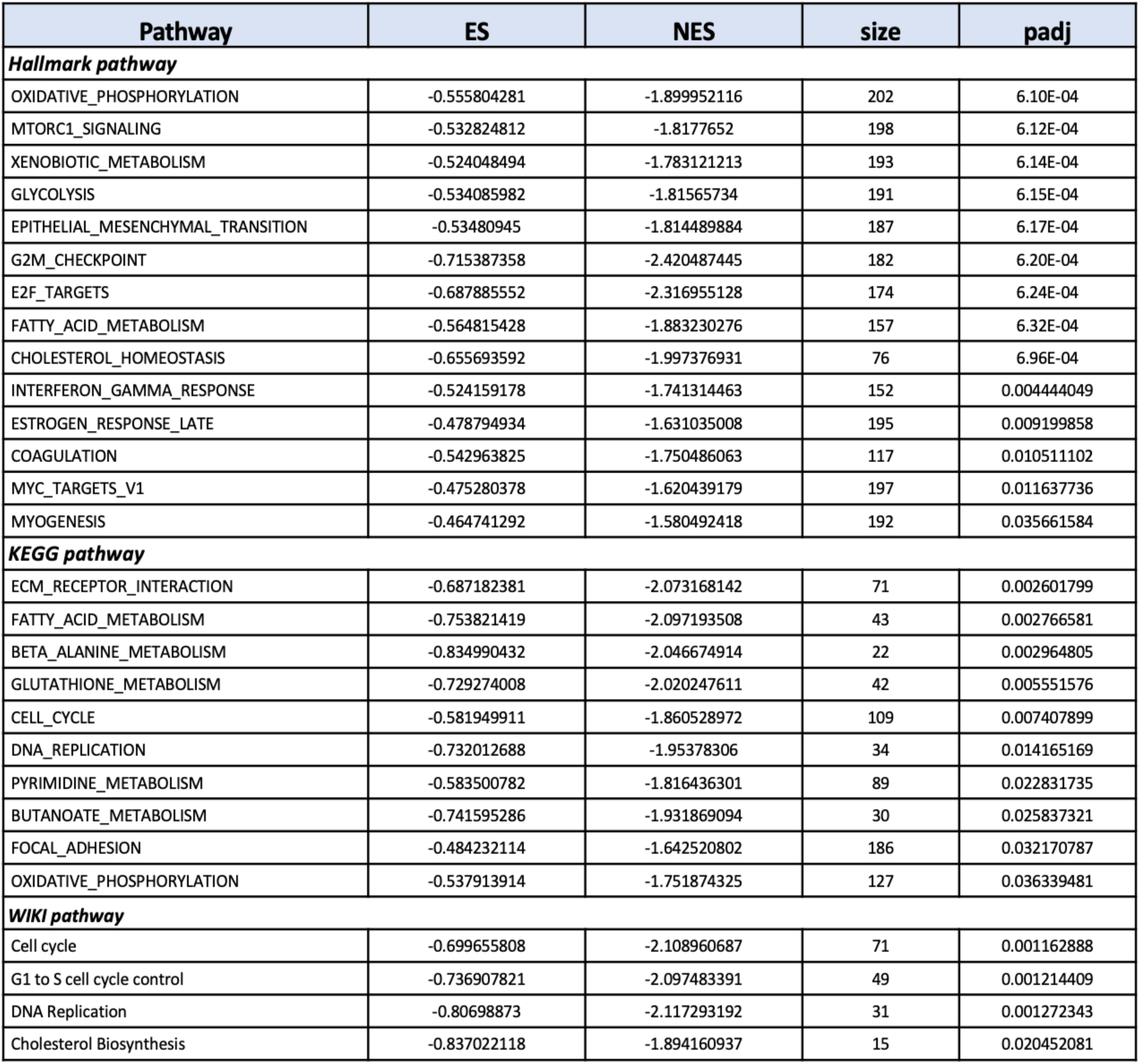
Significantly-changed pathways in the GSEA analysis of Hallmark, KEGG and Wiki pathways filtered by a Bonferroni correction P-value of 0.05. ES and NES indicate enrichment score and normalized enrichment score respectively. NES is generated through normalizing enrichment score to mean enrichment score of random samples. Size presents the numbers of genes contributing to each pathway.

**Figure 3.**
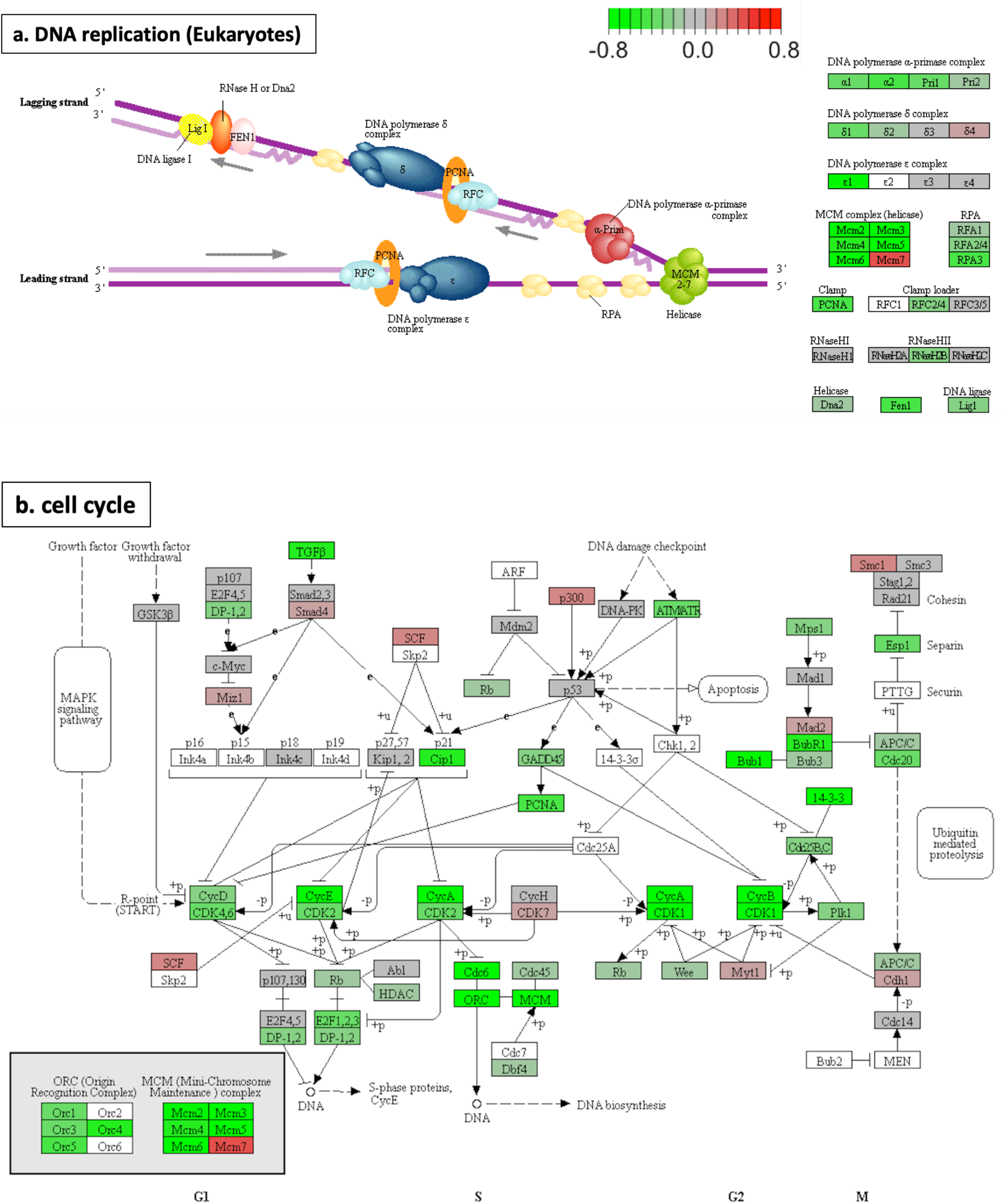
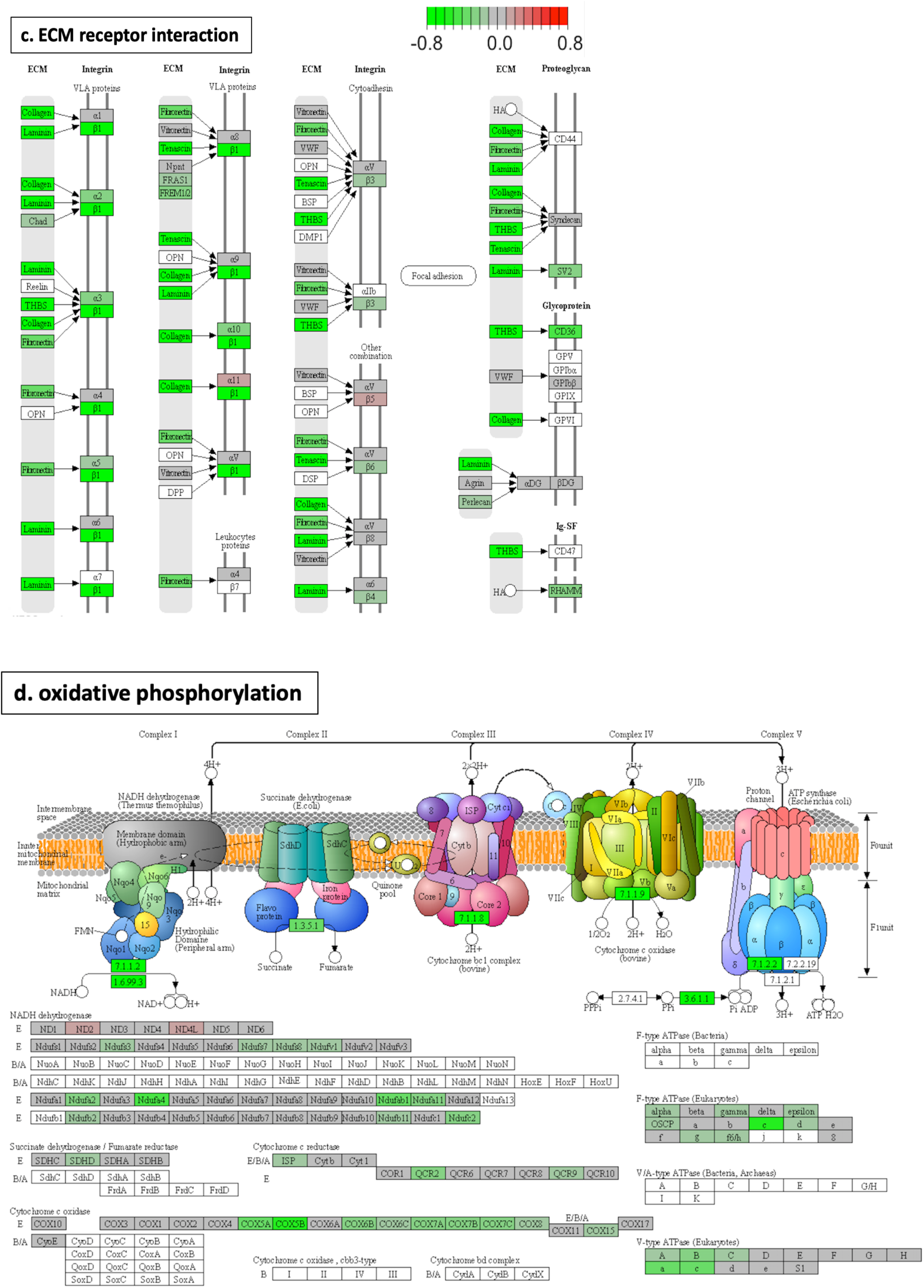
Diagrams for four significantly-changed KEGG pathways plotted using pathview [32]. a) DNA replication. b) cell cycle. c) ECM receptor interaction d) oxidative phosphorylation. Intensity of colour represents gene log_2_FC with downregulation in green and upregulation in red.

## Discussion

### No suitable transcriptome biomarkers identified for drug screening

Zebrafish larvae represent a powerful system for screening of chemical libraries in drug discovery [13]. However, the lack of consistency in transcriptome changes seen between heterozygous mutant *psen1*^*Q96_K97del*^ 7 dpf larvae compared to heterozygous mutant 6-month-old brains would appear to preclude the use of these mutant larvae to find drugs to suppress the transcriptome changes seen in the brains. Also, although the RNAs used for transcriptome analyses were extracted from clutches of larvae rather than individuals, the expression levels of the DE genes were, nevertheless, quite variable between clutches of the same genotype, either wild type or mutant (Supplementary data 2). We were able to identify significantly changed cellular pathways in common between the larvae and the brains. These related to oxidative phosphorylation, mitochondrial function and lysosomal acidification, indicating that similar stresses/biological effects caused by the presence of the *psen1*^*Q96_K97del*^ mutation likely exist during the entirety of zebrafish development from larvae to young adult. However, analysis of transcriptomes at this level would be unsuitable for the massively parallel screening of chemical libraries. Therefore, use of the *psen1*^*Q96_K97del*^ EOfAD-like mutation for discovery of AD-preventative drugs remains infeasible until a suitable biomarker can be identified.

### MCM complex dysregulation may drive DNA replication stress

Comparison of the transcriptomes of pools of 7 dpf heterozygous *psen1*^*Q96_K97del*^ mutant larvae to their wild type siblings revealed highly significant regulatory effects on genes involved with DNA replication and the cell cycle. These were identified by both Goseq analysis and GSEA. The majority of DE genes contributing to these two terms are related to the minichromosome maintenance (MCM) protein family. The eukaryotic MCM complex functions as a DNA helicase essential for DNA replication and cell division. The complex is comprised of the protein products of six genes, MCM2-7 (Figure3a, [45]). We observed a downregulation of zebrafish genes *mcm2, mcm3, mcm4, mcm5, mcm6* and *mcmbp* and an upregulation of *mcm7*. If this gene dysregulation phenomenon affects other vertebrates, including mammals, it may clarify a hitherto unexplained mutagenic effect of EOfAD mutations in, specifically, *PSEN1*. In 2002, Chan et al. [46] showed that forced expression of EOfAD mutant *PSEN1*, but not wild type *PSEN1*, increased the sensitivity of Rat pheochromocytoma (PC12) cells to DNA damage by etoposide. Responses included greater than normal increases in p53 protein levels and phosphorylation. In 2010, Michelsen et al. [47], studied the effects in mouse brains of transgenes expressing EOfAD mutant forms of the genes *APP*, or *PSEN1*, or both simultaneously. They observed an increase in the number of single-strand DNA breaks occurring in hippocampal granule cells of the dentate gyrus and hippocampal pyramidal cells in areas CA1/2 when brains expressed only the *PSEN1* mutant transgene. Interestingly, simultaneous expression of EOfAD mutant forms of both *APP* and *PSEN1* reduced the single stand break rate so that it was similar to that seen for only *APP* mutant transgene expression alone. In mice, reduced function of the *Mcm4* gene led to susceptibility to chromosome breaks induced by a DNA replication inhibitor, aphidicolin [48]. Notably, in 2011, Yurov et al. [42] suggested a DNA replication stress hypothesis of AD which proposes that replication stress caused by incomplete DNA replication leads to DNA damage or improper repair, subsequently resulting in the accumulation of genomic instabilities in AD brains. Genomic instabilities are associated with neurodegeneration in other aging-related diseases [49, 50]. Our observations, those of others (above), and Yurov et al.’s ideas suggest that associations between *PSEN1* functions and DNA integrity are an interesting area for further exploration and may give us greater insight into cellular stresses driving AD pathologies.

### Are crystallin genes upregulated due to disturbance of lysosomal acidification?

The discovery that DE genes were enriched under the GO term *structural constituent of eye lens*, focused our attention on the crystallin genes *crygm2d9, crygm2d17, cryba1l2*, and *cryaa*. Crystallin genes have not previously been linked with AD or *PSEN* functions. However, a paper by Valapala et al., 2013 [51] reported that the loss of the βA3/A1-crystallin gene in rat (*Cryba1*) reduces endolysosomal acidification, leading to reduced γ-secretase-mediated release of Notch intracellular domain (NICD) and impaired lysosomal-mediated degradation of Notch. Overexpression of *Cryba1* in a *Cryba1*^*-/-*^ knockout mouse apparently was able to rescue the deficient function of the vacuolar ATPase (v-ATPase) responsible for endolysosomal acidification. EOfAD mutations of human *PSEN1* are thought to reduce lysosomal acidification due to a requirement for the PSEN1 holoprotein for correct N-glycosylation of the v-ATPase subunit, V0a1 [52]. Our transcriptome analysis of the effect of heterozygosity for the *psen1*^*Q96_K97del*^ mutation on 6-month-old zebrafish brains also observed implied effects on lysosomal acidification [16]. While lysosomal acidification was not revealed as an affected GO in our analysis of 7 dpf heterozygous *psen1*^*Q96_K97del*^ larvae, we speculate that the upregulation of crystallin genes observed may be a homeostatic response to cope with disturbed lysosomal acidification.

Interestingly, Notch signaling has been seen to control expression of the protein products of the human *mcm* gene orthologues, *MCM2* and *MCM6* in cells where Notch signaling suppresses proliferation [53]. However, since Notch signaling was seen to downregulate these proteins, loss of Notch signaling due to increased lysosomal pH would not appear to explain the downregulation of *mcm2* and *mcm6* gene expression we observed in the heterozygous mutant zebrafish larvae.

### Does downregulation of ECM related genes increase the risk of AD?

The most significant KEGG pathway identified in GSEA was *ECM (extracellular matrix) receptor interaction*. Most genes in this pathway, including those encoding collagen, laminin, tenascin and thrombospondin, were significantly downregulated (Figure 3c). *ECM receptor interaction* is not currently a focus in AD studies, although altered regulation of these genes has previously been observed in some AD relevant research. In 2019, Kwart et al. [54] identified 1,515 overlapping DE genes from three human iPSC lines carrying EofAD mutations in *APP* and *PSEN1*. These DE genes were used to perform a enrichment analysis, and *ECM receptor interaction* was the second most statistically significant KEGG pathway identified. However, the relationships between EOfAD mutations and *ECM receptor interaction* have not been further explored. Another study by Conejero-Goldberg et al., 2014 [55] analysed human postmortem cortex and identified upregulation of ECM-related gene transcripts in carriers of the AD-protective ε2 allele of the gene APOE (i.e. APOE2). Based on this observation, they assumed increased ECM expression would reduce amyloid-β secretion or excitotoxicity. Thus, it appears that increased ECM gene expression is associated with decreased AD risk while EOfAD mutation-associated decrease of ECM gene expression promotes AD pathology. This may indicate the potential for ECM gene expression to act as an AD risk biomarker. More attention should be focused on ECM gene expression in future studies of AD.

### Mitochondrial dysfunction is an early effect of the *psen1*^*Q96_K97del*^ mutation

In addition to the pathways mentioned above, *fatty acid metabolism, oxidative phosphorylation* (*OX PHOS*) and *cholesterol biosynthesis* were identified in the GSEAs of KEGG and Wiki pathways. These pathways represent linked systems that control ATP production to meet cellular energy demand and in response to oxygen availability. Acetyl-CoA, as the starting point of *cholesterol biosynthesis*, is produced by oxidation reactions including oxidative decarboxylation of pyruvate and β-oxidation of fatty acids [56], and is a substrate in the tricarboxylic acid cycle (TCA cycle) to drive oxidative phosphorylation [57]. The AD brain is hypometabolic [58] and mitochondrial dysfunction is associated with oxidative stress in AD neuropathology through reduced ATP production [59]. Reductions in oxidative phosphorylation enzyme activities and functions have been identified in AD and other neurodegenerative processes [60]. A study by Manczak et al. [61] examined the expression of oxidative phosphorylation genes in AD patients and found downregulation of mitochondrial genes coding for electron transport chain (ETC) complex I, which is consistent with the gene expression in our mutant larvae (Figure 3d). However, Manczak et al. saw increased mRNA expression for components of complexes III and VI in contrast to our observations in mutant larvae. A more recent analysis by Mastroeni et al. [62] saw downregulation of nuclear-encoded ETC genes in AD but increased expression of these relative to age-matched controls in mild cognitive impairment (MCI). The downregulation of *fatty acid metabolism, oxidative phosphorylation* (*OX PHOS*) and *cholesterol biosynthesis* genes observed in mutant larvae indicates that the implied impairment of energy production by the EOfAD-like *psen1*^*Q96_K97del*^ mutation occurs early in life.

### Iron homeostasis

We have previously proposed that cellular iron dyshomeostasis may represent a unifying effect-in-common of the EOfAD mutations in *APP, PSEN1* and *PSEN2* [37] since APP was thought to stabilize the iron export protein FERROPORTIN [63], while endolysosomal acidification (affected by EOfAD mutations in *PSEN1* [52]) is important for import of iron into cells [64]. Recently, the role of APP in stabilization of FERROPORTIN has been challenged [65, 66]. However, it has been revealed that EOfAD mutations in APP, (like those in *PSEN1*), also affect acidification of the endolysosomal pathway [67] and so would be expected to affect cellular iron homeostasis. Our identification that the GO *iron ion transport* is affected in 7 dpf *psen1*^*Q96_K97del*^/+ larvae, particularly with downregulation of the *tfa* and *tfr1b* genes required for importation of iron, supports that EOfAD mutations in *PSEN1* disturb ferrous iron homeostasis. The fact that we have seen transcriptome evidence for such dyshomeostasis in 6-month-old *psen1*^*Q96_K97del*^/+ brains [16] but were unable to detect stabilization of mRNAs containing IREs in their 3’UTRs in 7 dpf larvae, suggests that any disruption of ferrous iron homeostasis begins subtly. Therefore, we propose that gene dysregulation promoting ferrous iron dyshomeostasis occurs by 7 dpf, but that the iron dyshomeostasis requires time to develop before it becomes apparent as ferrous iron dyshomeostasis in young adult *psen1*^*Q96_K97del*^/+ brains.

## Acknowledgements

The authors wish to thank Lachlan Warren William Baer for his technical assistance in construction of the GO plot.

