## Supplementary data 2 for "Transcriptome analyses of 7-day-old zebrafish larvae possessing a familial Alzheimer’s disease-like mutation in *psen1* indicate effects on oxidative phosphorylation, mcm functions, and iron homeostasis"

**Supplementary data 2: qPCR validation of RNA-seq data**

To support the accuracy and reliability of RNA-seq data, we performed relative standard curve quantitative PCRs (qPCRs) on four of the most statistically significantly DE genes with a relatively large fold-change in expression. Also, qPCRs were performed on three of the significantly changed genes identified in *GO iron ion transport* to confirm this reality. The cDNAs used in qPCRs were synthesized from the same RNA samples as were used in the RNA-seq.

**Method**

**Relative standard curve quantitative PCR**

1000ng of the total RNA from each sample remaining after RNA-seq was used to synthesise 20μL of first-strand cDNA by reverse transcription (SuperScript III kit, Invitrogen, Camarillo, California, USA). Standard curves were generated using a dilution series having 40ng, 20 ng, 10 ng and 5 ng of wild type cDNA per reaction. Each 25 μL qPCR reaction contained 20 ng of cDNA, 0.2 μM of each PCR primer and Power SYBR green master mix PCR solution (Applied Biosystems, Thermo Fisher Scientific Inc., Waltham, MA, USA). The qPCR was performed on an ABI 7000 Sequence Detection System (Applied Biosystems) using a 96-well plate. The amplification consisted of a holding stage and a cycling stage. The holding stage was performed at 50°C for 2 mins and then 95°C for 10mins, and the cycling stage had 40 cycles of 95°C for 15 s and 60°C for 1 min. Each reaction was conducted in triplicate. The quantities of amplified product in each reaction were determined from the standard curve. The mean value of three technical replicates was used to represent the quantity. The quantities of transcripts of the genes of interest were then calibrated to the quantities of transcripts of the house-keeping gene *rpl13* for relative quantification, enabling quantitative comparison between samples.

The gene expression measured by RNA-seq analysis was plotted as log_2_CPM (see the left half of the following figures in Results), while the qPCR results were plotted as quantities relative to *rpl13* (see the right half of the following figures in Results) with P-values calculated using a paired t-test.

**Primer Information**

| **Ensembl Gene ID** | **Primer name** | **Primer sequence** |
| --- | --- | --- |
| ***House-keeping gene*** | | |
| ENSDARG00000099380 | rpl13_F | 5’- CGCTAAGGACGGAGTGAACAAC-3’ |
|  | rpl13_R | 5’- CTCTCTTCTGCCAGTCTTTATGA-3’ |
| ***Statistically significantly DE genes*** | | |
| ENSDARG00000099511 | CABZ01034698.2_F | 5’- CTTATTGACGGAAAGATAGGAGAA-3’ |
|  | CABZ01034698.2_R | 5’- CTAGTCTGTGTCTAAATCTCTCATCGTG-3’ |
| ENSDARG00000103849 | mdh1ab_F | 5’- GCTTCCTCACAGGTGGCATT-3’ |
|  | mdh1ab_R | 5’- GGCAGTCTCGTCCTTCCCTT-3’ |
| ENSDARG00000077068 | si:ch211-11p18.6_F | 5’- TGCTGATACTCTGCTAAATCAACTG-3’ |
|  | si:ch211-11p18.6_R | 5’- CCTGTTCACTGCCACCCTGAG-3’ |
| ENSDARG00000093024 | si:ch211-213a13.2_F | 5’- CTCCAAAAGAGCCTTGTTAAATGC-3’ |
|  | si:ch211-213a13.2_R | 5’- TTACTGAGCCCCAATATCTCCAA-3’ |
| ***Genes identified in the GO* iron ion transport** | | |
| ENSDARG00000016771 | tfa_F | 5’- GCAATCCCAGAGAGTGAGAGG-3’ |
|  | tfa_R | 5’- CTGCTTCAGGACATCATAAACAT-3’ |
| ENSDARG00000077372 | tfr1b_F | 5’- CCGTATTATGAAGGTTGAGCA-3’ |
|  | tfr1b_R | 5’- CAGGTCTGTGGCGGTTTGC-3’ |
| ENSDARG00000094210 | fthl31_F | 5’- ATAGTAAAACCTGCCGAGATGGA-3’ |
|  | fthl31_R | 5’- TAAAATAGTGAGCCATGGAGGTG-3’ |

**Results**

**Validation of statistically significantly DE genes**

QPCRs were performed on transcripts of the genes *CABZ01034698.2, mdh1ab, si:ch211-11p18.6* and *si_ch211-213a13.2.* The relative quantifications of *CABZ01034698.2, mdh1ab* and *si:ch211-11p18.6* indicated significant differences (P-value < 0.05) between wild type and heterozygous mutant larvae with the same regulation direction as observed in RNA-seq. However, the relative quantification of *si_ch211-213a13.2* was different from the RNA-seq result, probably due to mis-mapping or misidentification of reads in the RNA-seq analysis. Therefore, three of four statistically significantly DE genes identified in RNA-seq were validated by qPCR, supporting the accuracy of most RNA-seq results.

***CABZ01034698.2***

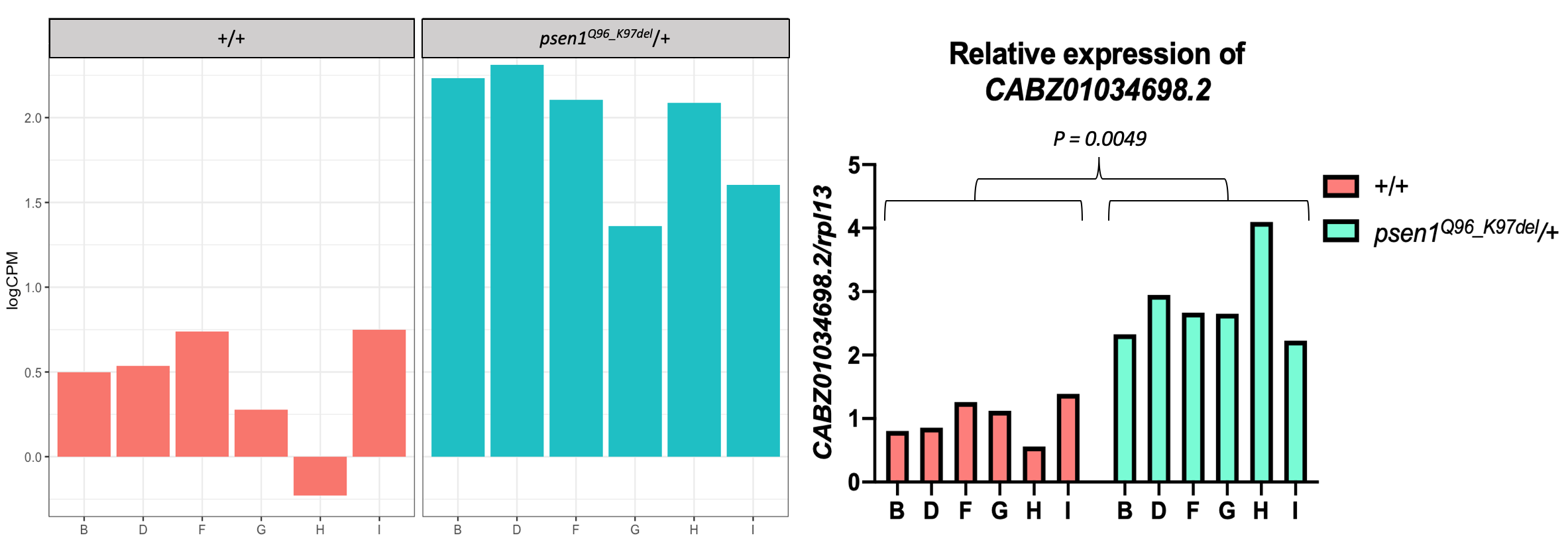

***mdh1ab***

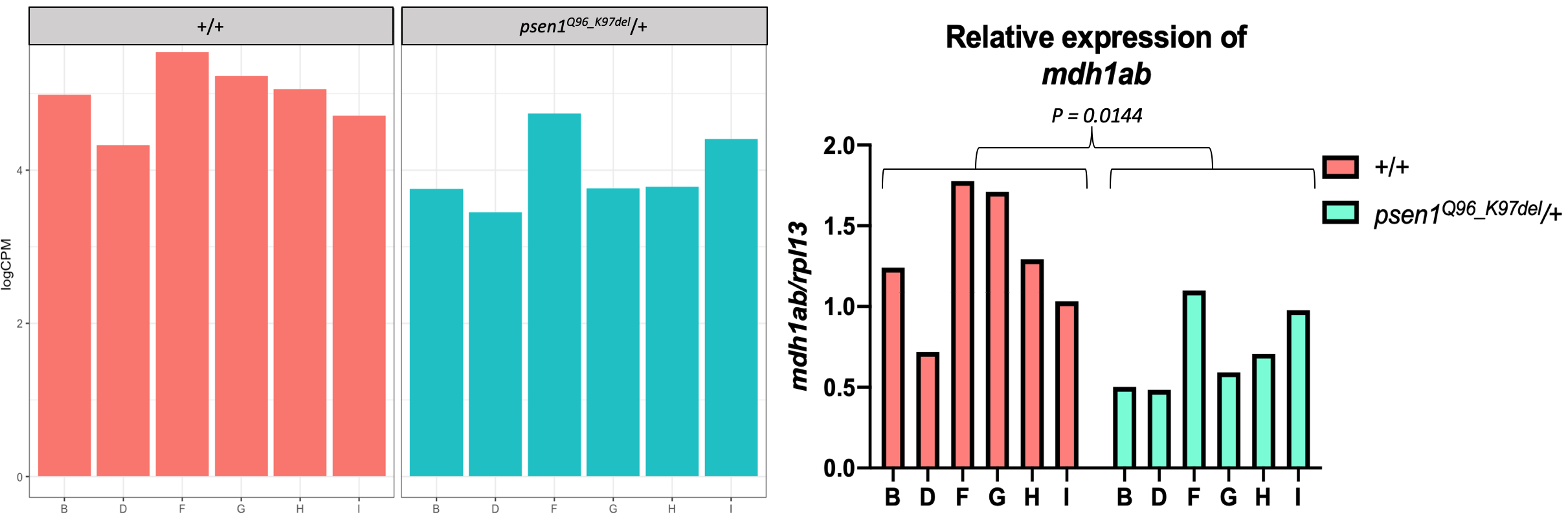

***si:ch211-11p18.6***

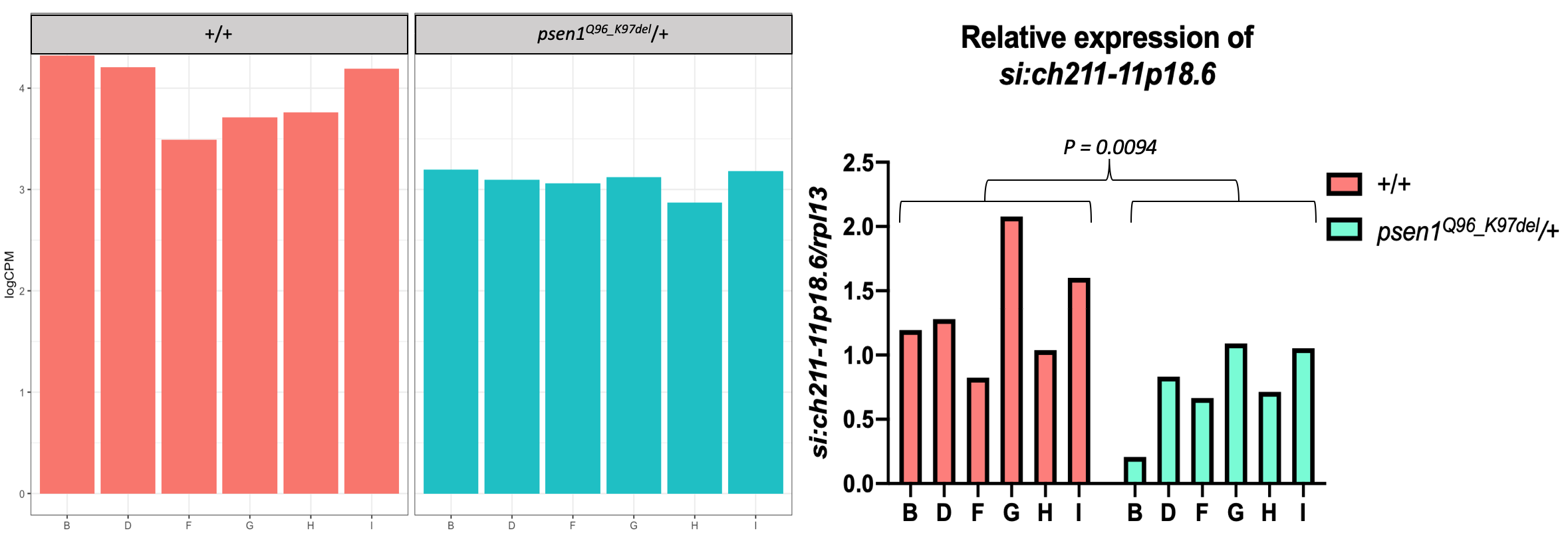

***si_ch211-213a13.2***

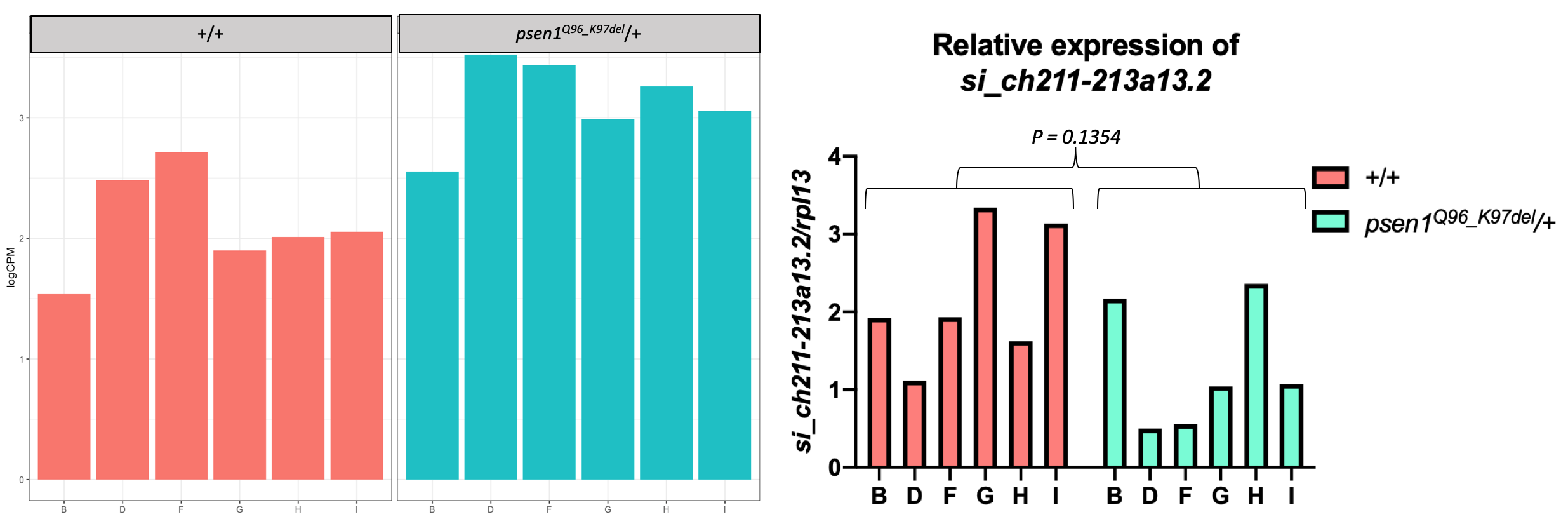

**Validation of the genes identified in the GO *iron ion transport***

The same qPCR approach as above was applied to the genes *tfa*, *tfr1b* and *fthl31* identified in the GO *iron ion transport*. Only the difference in relative expression level of *fthl31* validated the RNA-seq result at the statistical significance threshold chosen (P-value < 0.05) in a paired t-test. However, although the differences in relative expression levels for *tfa* and *tfr1b* did not show statistical significance, the relative expression level in each sample generally corresponded with the relative log_2_CPM level in RNA-seq analysis, indicating that the qPCR results supported the reality of the RNA-seq results. Also, detection of statistically significantly DE genes by RNA-seq included all the detected genes of the transcriptome in estimating variance, while qPCR only examined variation in the gene of interest across samples and so was less sensitive in detecting statistical significance.

***tfa***

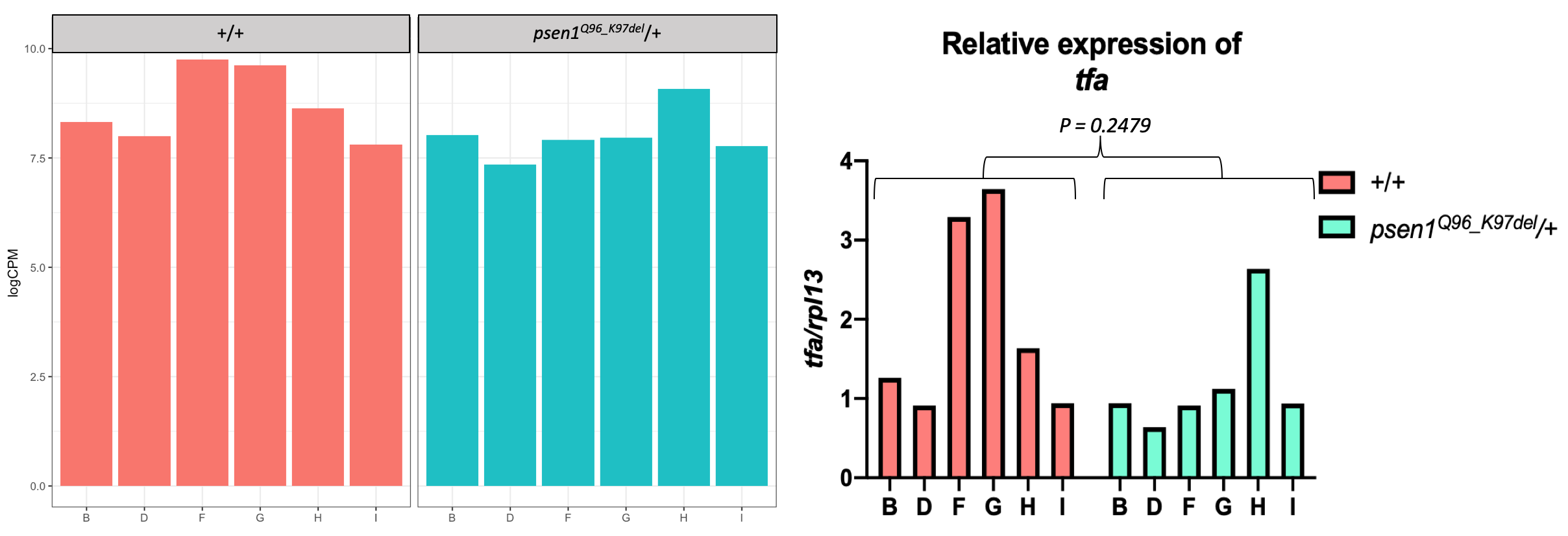

***tfr1b***

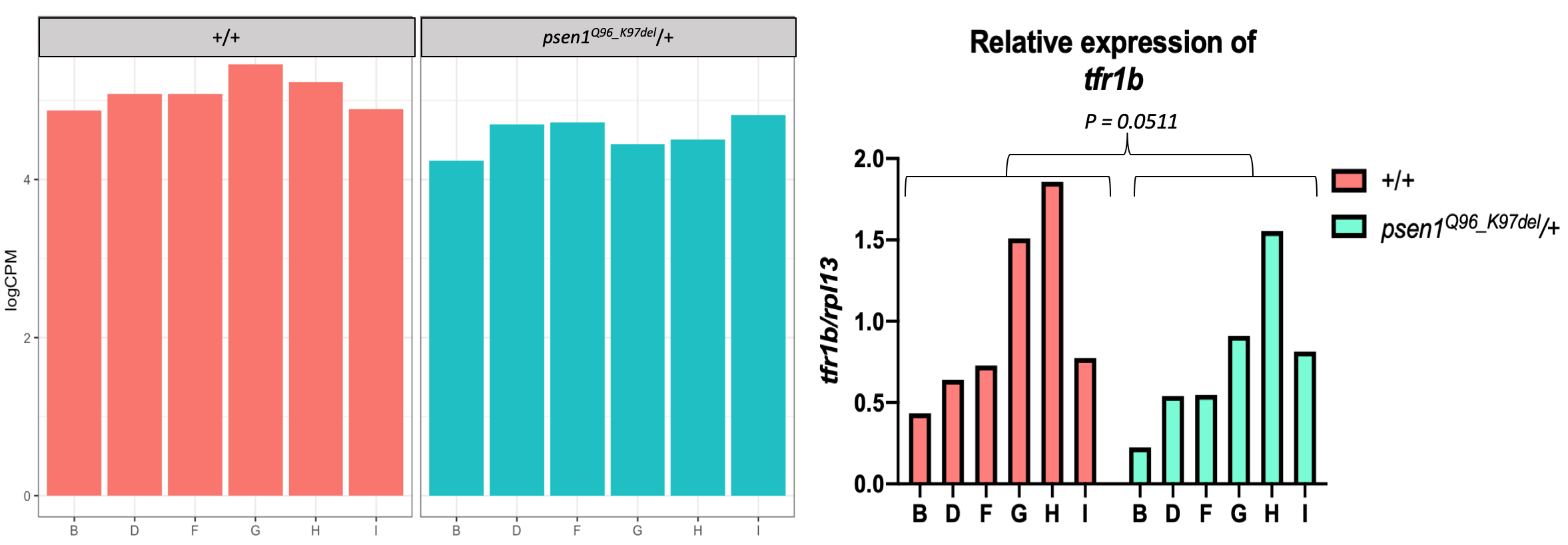

***fthl31***

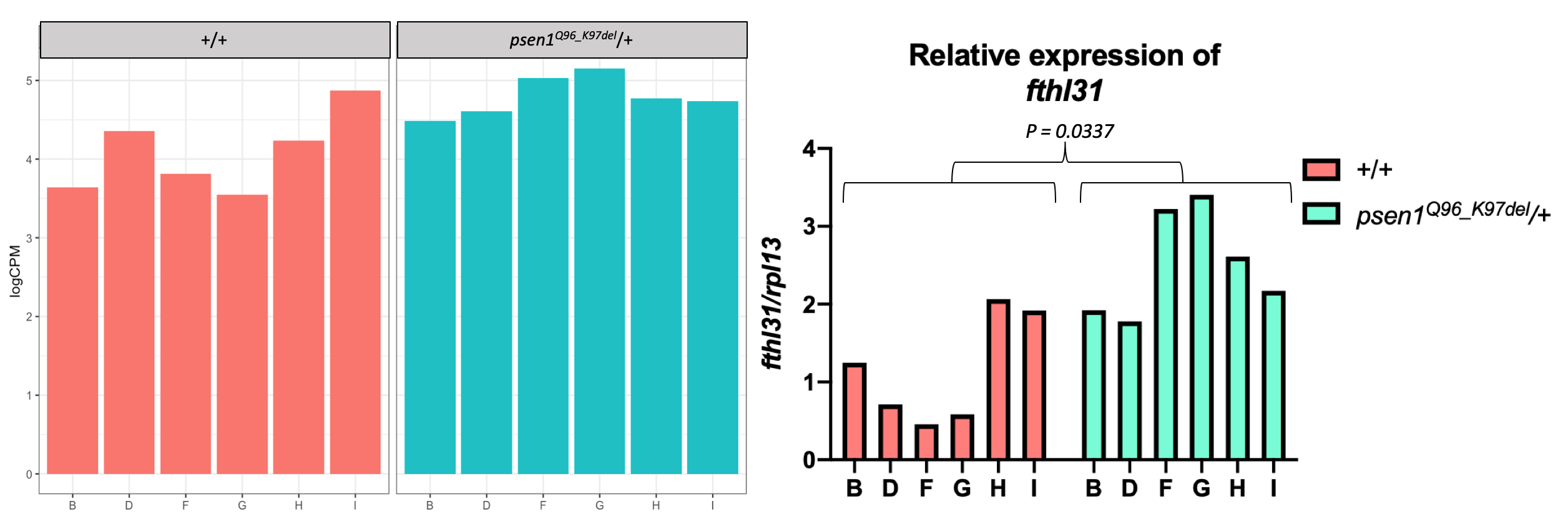

Both the RNA-seq plots and the qPCR plots display obvious variability in gene expression within clutches of larvae of the same genotype. For example, although the expression levels of *fthl31* in wild-type clutches H and I were similar to the expression levels in mutant clutches B and D, when analyzed in pairs, the expression levels of *fthl31* in the mutants were obviously higher than in the wild-type clutches. In this sense, our paired-mating strategy (Figure 1 of the paper) aided in DE gene identification.
