## Supplementary data 3 for "Transcriptome analyses of 7-day-old zebrafish larvae possessing a familial Alzheimer’s disease-like mutation in *psen1* indicate effects on oxidative phosphorylation, mcm functions, and iron homeostasis"

**Supplementary data 3: KEGG pathway diagrams**

Ten significantly-changed KEGG pathways were identified by GSEA. The KEGG diagrams of *DNA replication, cell cycle, ECM receptor interaction* and *oxidative phosphorylation* are shown in the main paper. The diagrams of the other 6 significantly changed KEGG pathways are listed below. Intensity of colour in diagrams represents gene log_2_FC ranging from -0.8 to 0.8 with downregulation in green and upregulation in red.

**Focal adhesion**

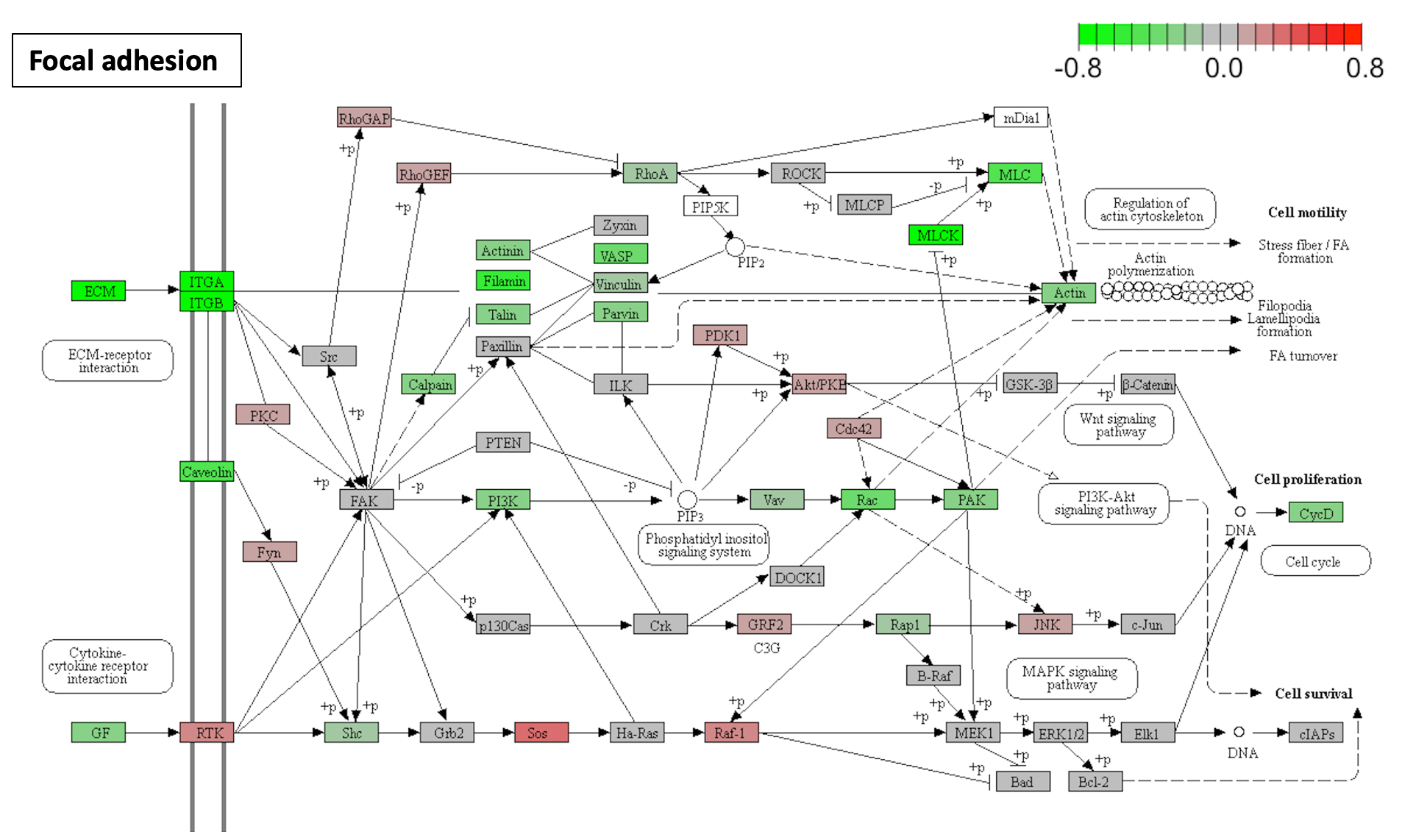

**Fatty acid metabolism**

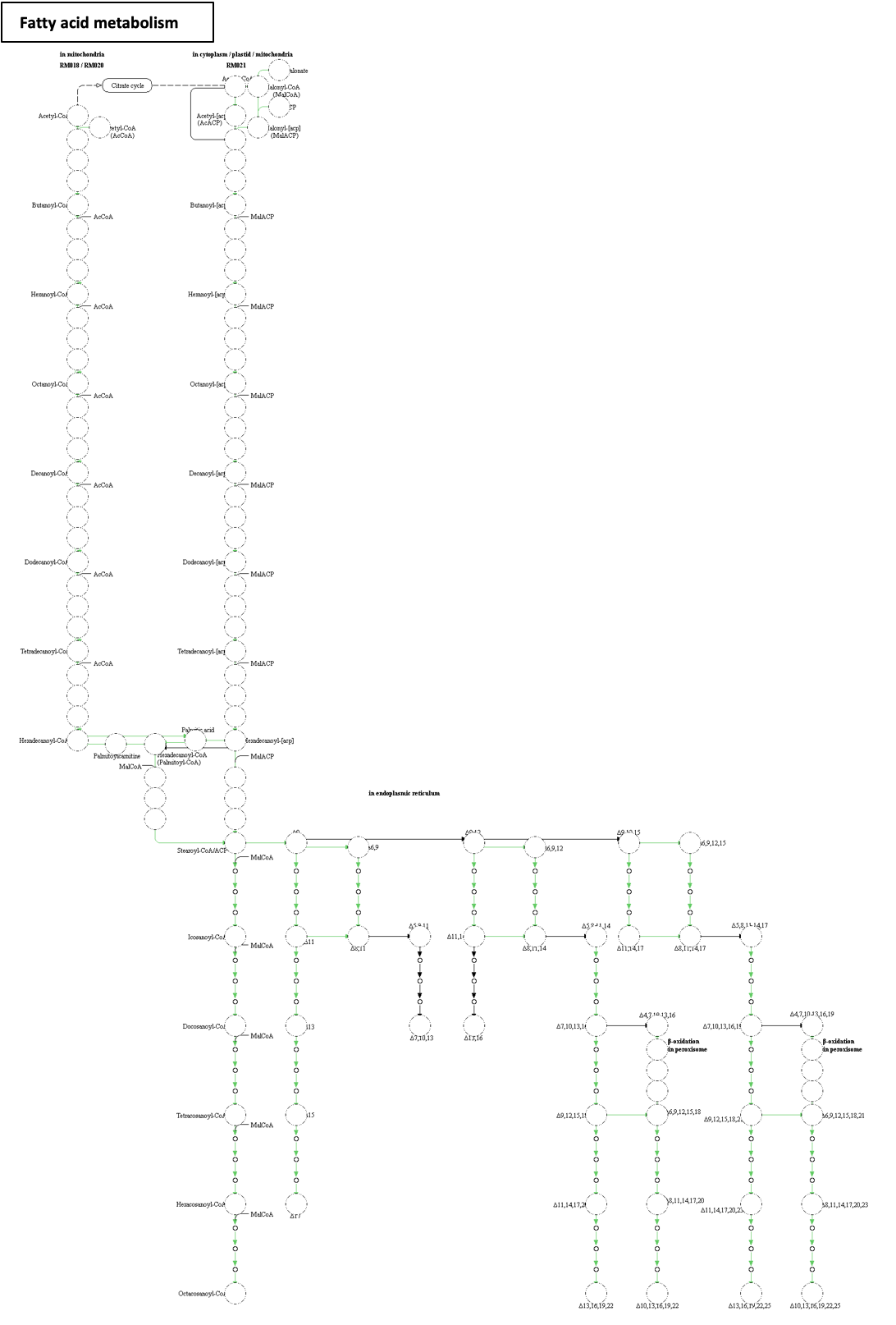

**Beta-Alanine metabolism**

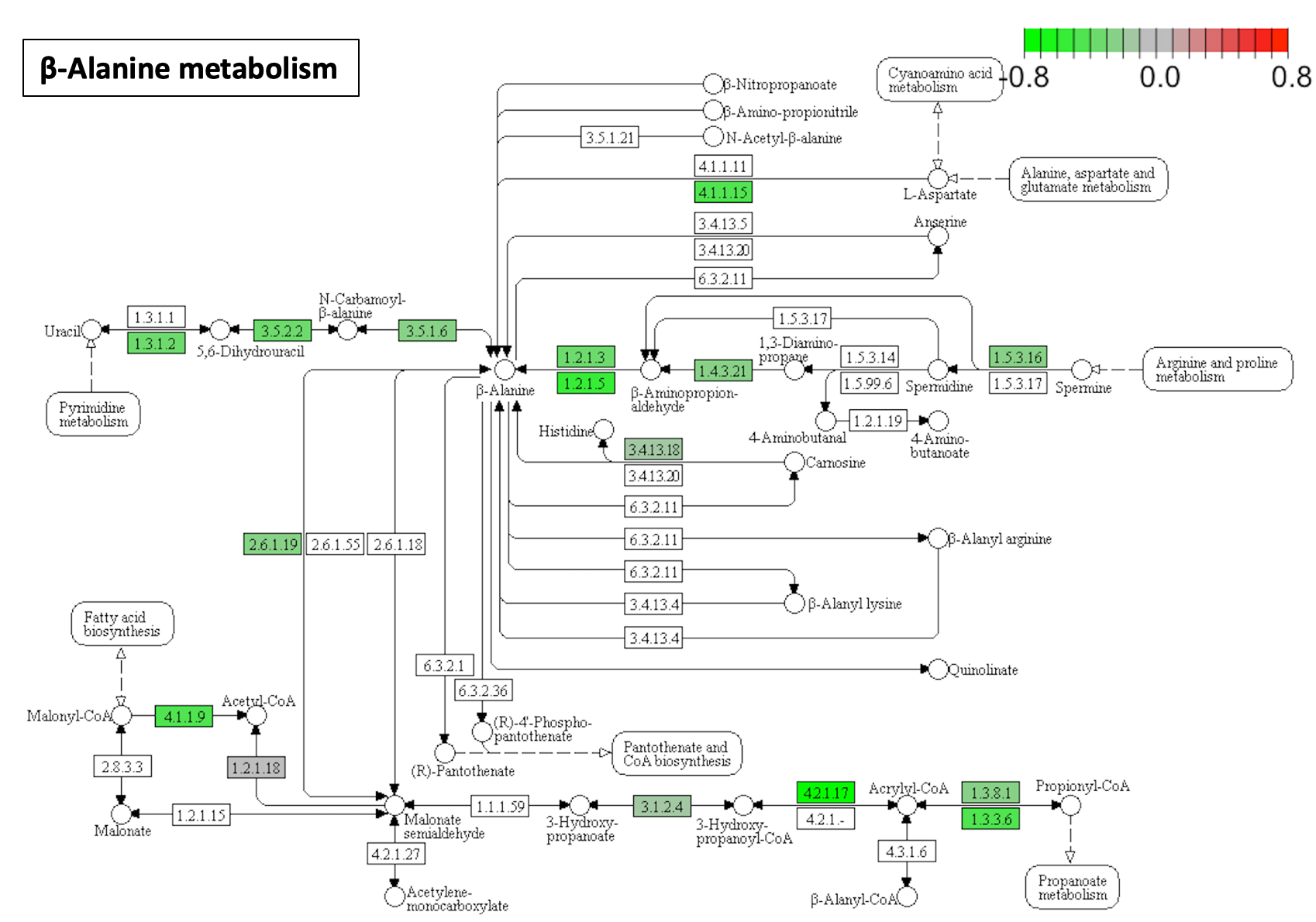

**Glutathione metabolism**

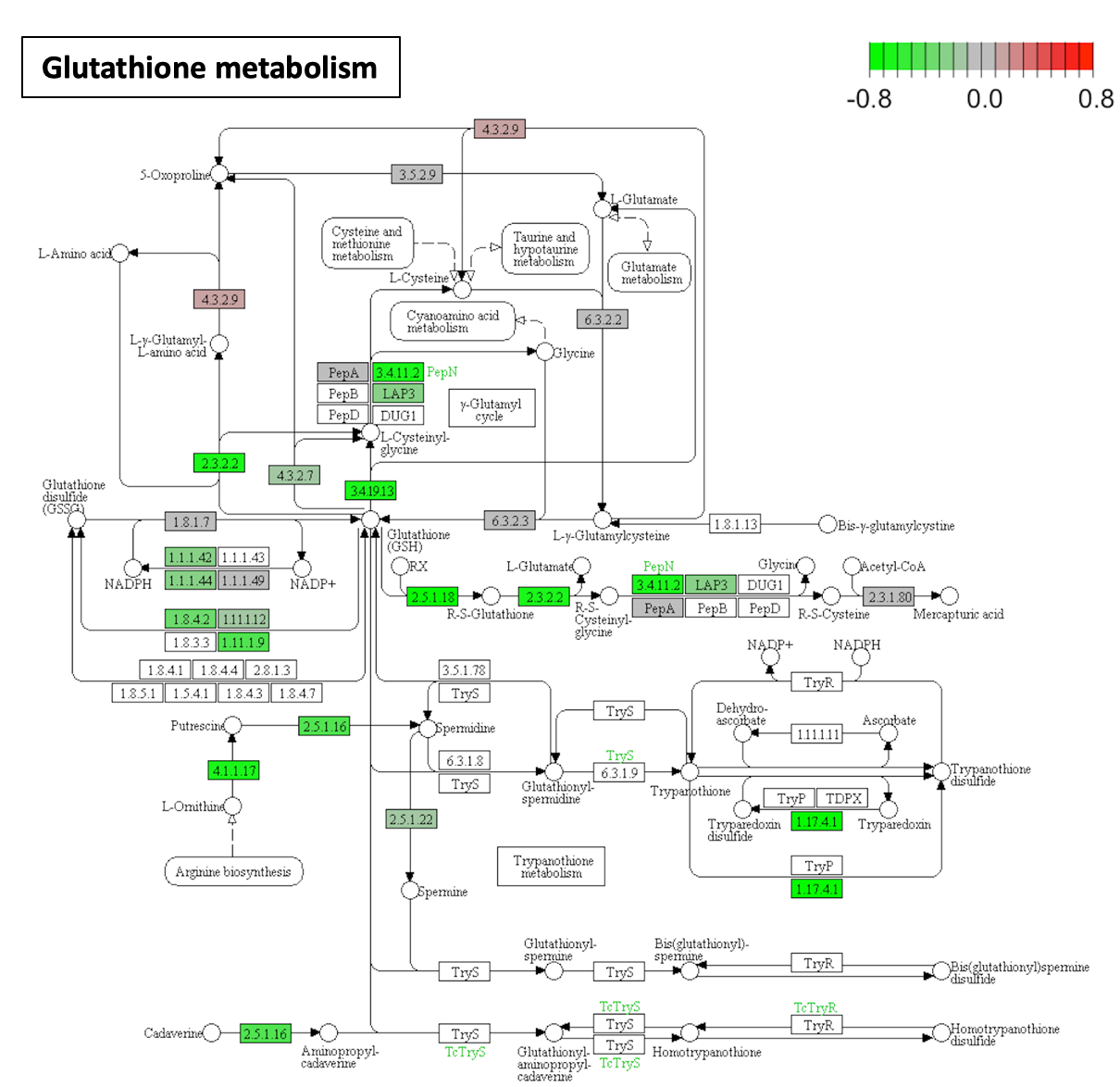

**Pyrimidine metabolism**

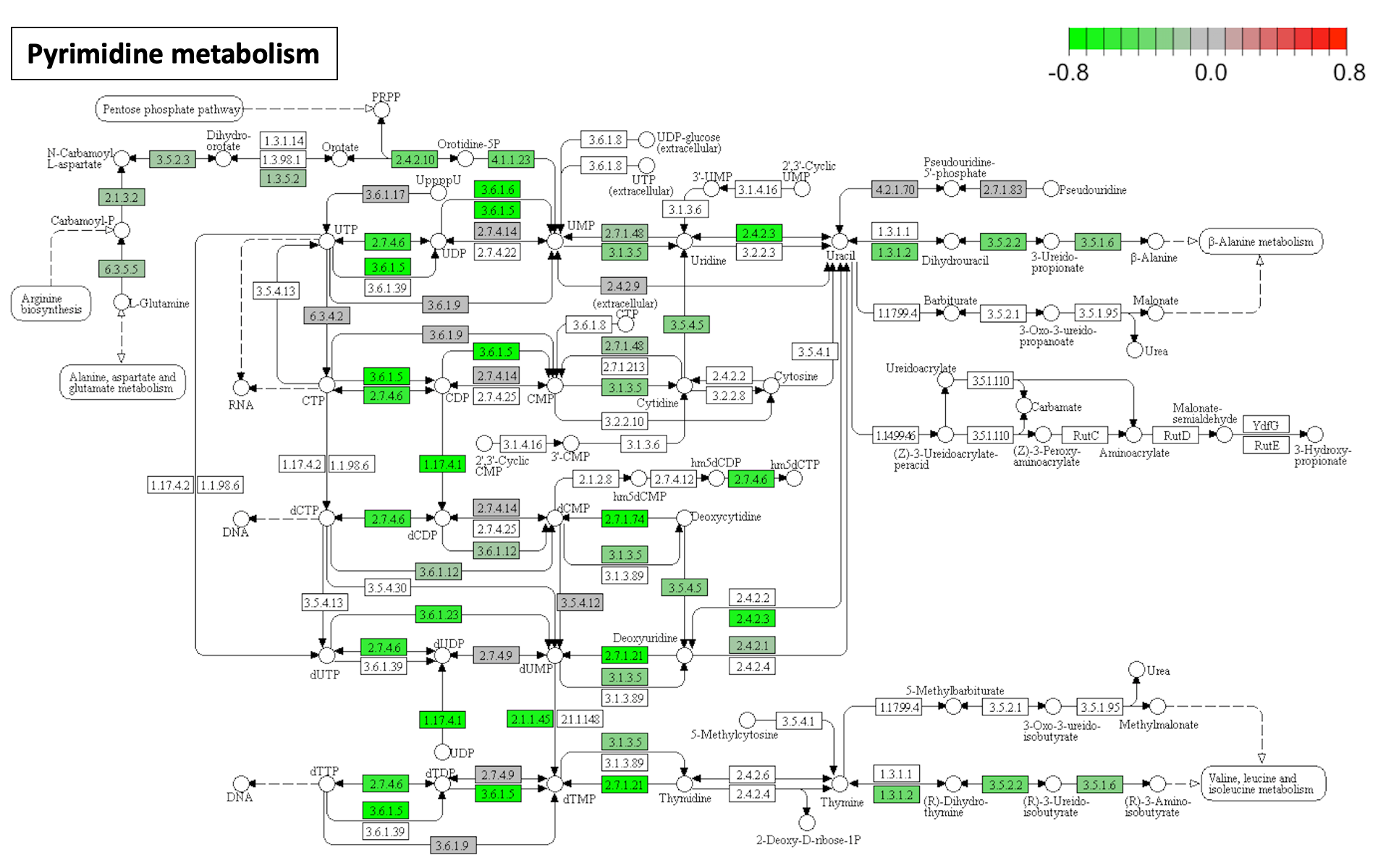

**Butanoate metabolism**

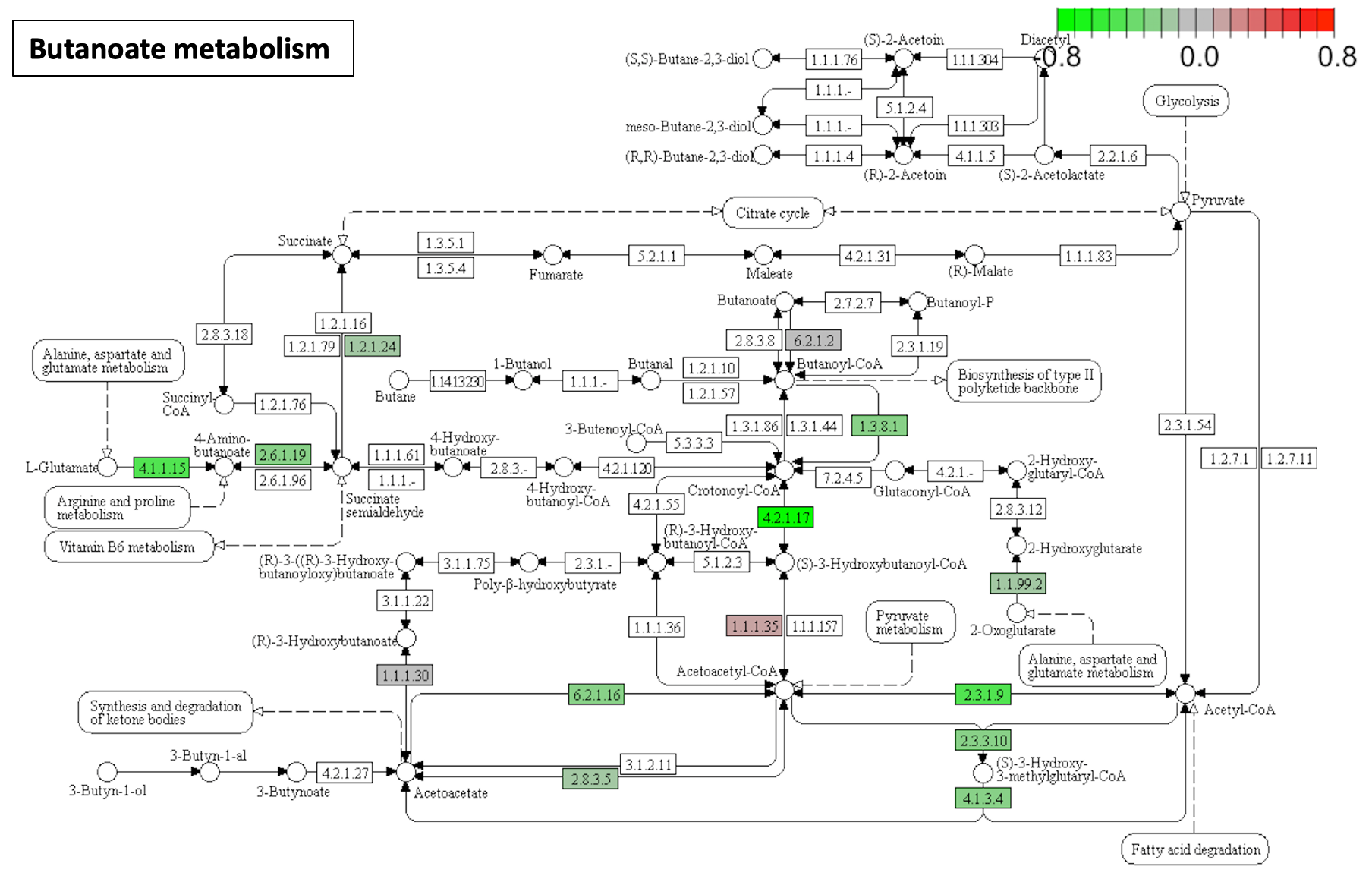
