## Supplementary data 4 for "Transcriptome analyses of 7-day-old zebrafish larvae possessing a familial Alzheimer’s disease-like mutation in *psen1* indicate effects on oxidative phosphorylation, mcm functions, and iron homeostasis"

**Supplementary data 4: IRE enrichment analysis**

The GO *iron ion transport* was identified as significantly changed in heterozygous *psen1^Q96_K97del^* mutant larvae, indicating the influence of the *psen1^Q96_K97del^* mutation on iron homeostasis. Here, in order to detect whether ferrous iron dyshomeostasis effects are evident in the transcriptome data, we performed enrichment analysis of the gene sets containing IREs (iron-responsive elements) in either their 5’ or 3’ UTRs as defined by Hin et al., 2020 [1] using Goseq analysis (Supplementary data 4, Table 1) and GSEA (Supplementary data 4, Table 2). Genes containing high confidence IRE predictions were also selected as independent categories for enrichment analysis.

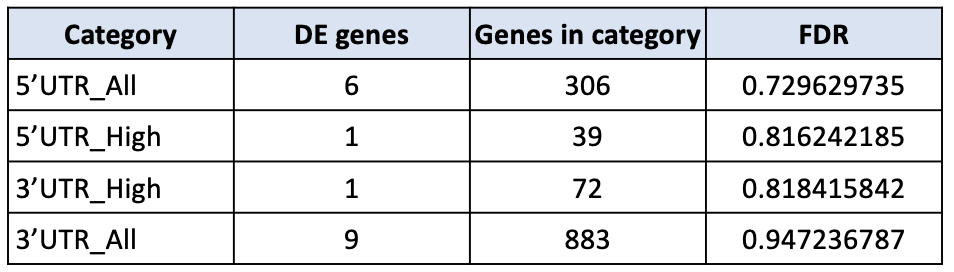

**Supplementary data 4, Table 1.** Enrichment analysis of the sets of genes containing IREs using Goseq analysis. “All” indicates all genes with such predicted IREs, while “High” contains only genes with such IREs predicted with high confidences (see [1]). FDR, false discovery rate.

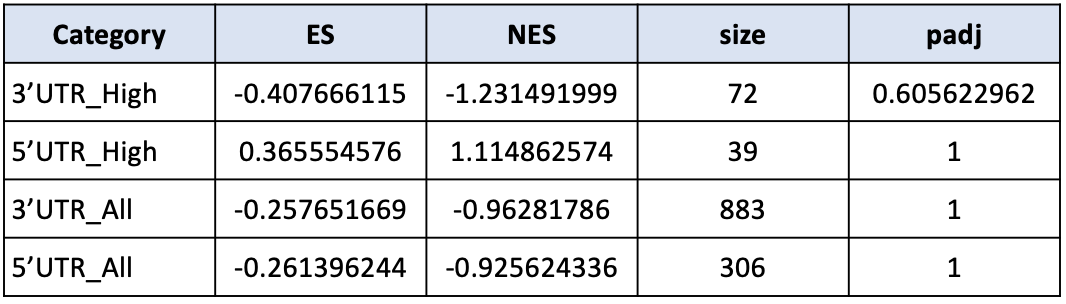

**Supplementary data 4, Table 2.** Enrichment analysis of the sets of genes containing IREs using GSEA. ES and NES indicate enrichment score and normalized enrichment score respectively. “Size” presents the numbers of genes contributing to each category. padj, adjusted P-value.

In both Goseq analysis and GSEA, none of the gene sets showed significant enrichment (P-value < 0.05), indicating that the ferrous iron dyshomeostasis identified in young adult *psen1^Q96_K97del^*/+ brains was not detectable by this method at the transcriptome level in 7 dpf zebrafish larvae.
